# Recruitment of Fpt1 to tRNA genes requires TFIIIB and the N-terminal TPR array of TFIIIC subunit τ131

**DOI:** 10.64898/2025.12.24.695929

**Authors:** Maria Elize van Breugel, Wolfram Seifert-Davila, Marianne J. Bakker, Joseph V. W. Meeussen, Ilse A. Hoogvliet, Mireia Novell Cardona, Tibor van Welsem, Danique Ammerlaan, Florence Baudin, Patrick Celie, Tineke L. Lenstra, Christoph W. Müller, Fred van Leeuwen

## Abstract

Transfer-RNA genes (tDNAs) in budding yeast recruit varying amounts of Fpt1, a regulator of RNA polymerase III (RNAPIII) occupancy. Fpt1 occupancy resembles that of the general transcription factor TFIIIC but how Fpt1 is recruited to tDNAs remains unclear. Here we show that both TFIIIB and TFIIIC are required for Fpt1 binding under active as well as repressive RNAPIII conditions. Depletion of TFIIIB reduced Fpt1 occupancy without affecting TFIIIC. In contrast, TFIIIC depletion led to reduced Fpt1 and a gene-specific reduction in TFIIIB occupancy. Moreover, upon depletion of TFIIIC, Fpt1 and TFIIIB were lost to different extents. We identified the C-terminal intrinsically disordered region of Fpt1 as critical for increased Fpt1 binding under repressive conditions. Within this region, a short α-helix was predicted to interact with the N-terminal tetratricopeptide repeat array of the TFIIIC subunit τ131, a region also known to interact with TFIIIB. Deletion of this α-helix abrogated stress-induced Fpt1 recruitment to tDNAs, as did mutations in the predicted interaction surface of τ131, while having a milder effect on TFIIIB occupancy. Together, these findings uncovered a dual and dynamic mechanism of Fpt1 recruitment to tDNAs with independent contributions of TFIIIB and TFIIIC.

## INTRODUCTION

RNA polymerase III (RNAPIII) is a large multi-subunit enzyme that transcribes genes encoding small noncoding RNAs. The majority of the RNAPIII repertoire comprises transfer-RNA (tRNA) genes (tDNAs), with approximately 275 and 400 copies in the yeast and human genomes, respectively [1]. RNAPIII-transcribed genes are classified into three categories based on promoter architecture; tRNA genes belong to Type II genes, characterized by internal promoter elements known as the A-box and B-box [2–5]. These conserved elements serve as binding sites for the six-subunit assembly factor TFIIIC [6,7]. TFIIIC consists of two submodules, τA (in *Saccharomyces cerevisiae* composed of subunits τ131, τ95, τ55, and the C-terminal region of τ138) and τB (τ91, τ60, and the N-terminal region of τ138) that bind the A- and B-box respectively [8]. Given that TFIIIC binds the A-box with micromolar affinity and the B-box with nanomolar affinity, its recruitment is primarily mediated by the B-box [9–11]. The flexible linker in τ138 allows TFIIIC to engage with A- and B-boxes positioned at varying distances [12].

Once TFIIIC is correctly positioned on the tRNA gene, it facilitates the stepwise recruitment of TFIIIB (in yeast composed of Brf1, Bdp1 and TBP) upstream of the transcription start site [13]. First, Brf1 binds to the N-terminal moiety of τ131 including the tetratricopeptide repeat (TPR) array, which promotes a conformational change in TFIIIC that allows TBP and Bdp1 recruitment [14–16]. It is hypothesized that once TFIIIB is assembled upstream of the tRNA gene, TFIIIC is dispensable for correct RNAPIII positioning and transcription [10,13,17,18]. Whether TFIIIC is fully released from the chromatin during RNAPIII elongation is topic of ongoing debate. As opposed to its activating role by recruitment of TFIIIB, TFIIIC is also proposed to act as a transient barrier of RNAPIII transcription since its occupancy at tRNA genes is increased under repressive conditions [19–24]. Under these conditions, the general RNAPIII-repressor Maf1 is de-phosphorylated, which leads to its translocation to the cell nucleus. Here, Maf1 directly interacts with RNAPIII and TFIIIB to prevent de-novo assembly onto the DNA [25–27]. This causes a decrease in RNAPIII occupancy (∼65-75% reduction), as well as a drop in TFIIIB (∼45-50% reduction), although to a lesser extent [19,21,22].

Maf1 is not a component of the tDNA chromatin itself and is considered a trans-factor. A recently identified negative regulator of RNAPIII, Fpt1, is suggested to act in parallel to Maf1 directly at the tDNA chromatin level [19]. Unlike Maf1, Fpt1 is constitutively present in the nucleus, and already bound to tDNAs in conditions of active transcription. However, the level of Fpt1 binding varies across tRNA genes, with higher levels found at stress-responsive tRNA genes and lower levels at housekeeping tRNA genes. Under repressive conditions, the occupancy of Fpt1 at tRNA genes is increased. In the absence of Fpt1, the eviction of RNAPIII and TFIIIB from tRNA genes is partially compromised by mechanisms that are currently unknown [19]. To ultimately uncover such mechanisms, in this study we investigated how Fpt1 is recruited to tDNAs. This recruitment varies across tDNAs (even between genes with identical body sequences) suggesting that gene context and DNA sequence influence Fpt1 recruitment [19]. While these factors may influence Fpt1 recruitment, protein interactions are also likely to play a major role. Here, we focus on how Fpt1 interacts with the protein complexes at tRNA genes. Several lines of evidence suggest that Fpt1 contacts tDNAs through interaction with the general transcription factors TFIIIB and TFIIIC. The overlapping ChIP-exo patterns between Fpt1 and individual subunits of TFIIIB and TFIIIC suggest close physical proximity of these factors [19]. In addition, conditional depletion of TFIIIB and TFIIIC in conditions of active transcription abolishes Fpt1 binding to tRNA genes, showing that these factors are necessary for Fpt1 recruitment [19]. Notably, Fpt1 ChIP-exo patterns and dynamics upon nutrient perturbation align more closely with TFIIIC than TFIIIB [19]. This raises the question of how Fpt1 depends on both TFIIIC and TFIIIB, despite their opposing dynamics under nutrient stress. Gaining more insight into how Fpt1 is recruited to tRNA genes in changing physiological conditions will benefit our understanding of the molecular mechanism by which Fpt1 regulates RNAPIII assembly.

Here, we combine conditional protein depletion, mutagenesis, and structure-based interaction prediction strategies to investigate the dynamic interaction of Fpt1 with the core RNAPIII transcription machinery. Protein depletion experiments show that for its association with tDNAs, Fpt1 relies on both TFIIIB and TFIIIC independently, regardless of nutrient availability. Mutational analysis revealed a stress-responsive region within Fpt1 that is essential for its increased tDNA occupancy under repressive conditions. This domain is predicted to interact with the TFIIIC subunit τ131. Notably, mutations in the predicted Fpt1-interacting region in τ131 impaired the stress-induced increase in Fpt1 occupancy at tDNAs, independently of TFIIIB. Together, our findings suggest a dual and condition-dependent recruitment mechanism whereby Fpt1 relies on both TFIIIB and TFIIIC to interact with tRNA genes.

## MATERIALS AND METHODS

### Yeast strains and plasmids

All strains, plasmids and oligos used in this study are listed in **Supplemental Table S1**. To construct yeast strains, haploid yeast cells (*Saccharomyces cerevisiae*) were transformed using the standard LiAc/ssDNA method [28]. The parental strain was BY4741 unless otherwise specified.

Strains NKI5650 and NKI5651 were isolated from the NKI5616 Epi-Decoder library [19]. NKI5754 was made using plasmid pFvL029 and the *Fpt1(Δ254-353)-TAP* oligos. Subsequently, NKI2714 was made from NKI5754 with a repair template from plasmid pTW183 and oligos *Fpt1(Δ254-353)-BPSV40* and CRISPR-Cas9 plasmid pMNC2. For NKI5786 and NKI5803 (with parental strain NKI5632), the CRISPR-Cas9 containing plasmids pMvB16 and pIAH001 were used in combination with *Repair template Fpt1Δ290-314* and *Repair template Fpt1Δ2-253* respectively. NKI5811 originated from NKI5803 and was made with plasmid pYM-N15 and oligos *Fpt1(Δ2-253)-TAP-TDH3*.

NKI5755 was made using plasmid pKT127 and oligos *Fpt1(Δ254-353)-GFP*. Subsequently, NKI2713 was made from NKI5755 with a repair template from plasmid pTW183 and oligos *Fpt1(Δ254-353)-BPSV40* and CRISPR-Cas9 plasmid pMNC1. For NKI5757 and NKI5758, NKI5755 and NKI2713 were transformed respectively with PacI digested pTL306 to insert a fluorescent coat protein and an URA3 selection marker at the *URA3* locus. NKI5787, NKI5791 and NKI5796 (with parental strain NKI5756) were made with CRISPR-Cas9 plasmid pMvB17 and *Repair template Fpt1 NLS(331-339)*, *Repair template Fpt1 NLS(319-353)*, and *Repair template Fpt1 NLS(324-339)* respectively. NKI5788 and NKI5802 (with parental strain NKI5756) were made using CRISPR-Cas9 plasmids pMvB16 and pIAH001 and *Repair template Fpt1Δ290-314* and *Repair template Fpt1Δ2-253* respectively.

NKI5774, NKI5775, NKI5776, and NKI5792 originate from parental strain BY4741-AA [29] and were made using plasmid pFvL029 and oligos *Brf1-TAP*, *Rpo31-TAP*, *Tfc3-TAP,* and *Tfc1-TAP* respectively. NKI5779 and NKI5781 originate from NKI5774 and NKI5776 respectively and were made using plasmid pTL100 and oligos *Tfc3-FRB* and *Brf1-FRB* respectively. For strain NKI5806, a repair template from pBS1761 using oligos *TAP-Tfc4* was used in combination with CRISPR-Cas9 plasmid pMvB18. Similarly, NKI5809 was made with pMvB18 and a repair template from pFvL160 using oligos *HA-Tfc4*. NKI5810 (with parental strain BY4741-AA) was constructed using plasmid pMvB18 and a repair template from pTL100 with oligos *FRB-Tfc4*. Subsequently, NKI5814 and NKI5818 were made from NKI5810 using plasmid pFvL029 and oligos *Fpt1-TAP* and *Brf1-TAP* respectively.

NKI7100 was made in the BY4741-AA background with a PCR product from genomic DNA of YTL1446 with the *Tfc3-FRB-V5* oligos. For NKI7111, *Brf1-FRB-V5* oligos were used to amplify the FRB-GFP-3xV5-HPHMX tag from the *Tfc3-FRB-3xV5-HPHMX template DNA* for integration in BY4741-AA.

For plasmid construction, PCR products from plasmids were digested with Dpn1 (1h at 37°C) to get rid of template DNA. Digested plasmids were gel purified using the QIAquick Gel Extraction Kit (Cat#28706). CRISPR-Cas9 plasmids pIAH001, pMNC1, pMNC2, pMvB16, pMvB17 and pMvB18 were constructed by cloning their respective sgRNAs into pML104-HygMx4 or pML104-KanMx4 using the BclI and SwaI restriction sites. Plasmid pTW183 was constructed Gibson Assembly (NEB, Cat#E2621L) of SalI digested pVZ1 and a PCR fragment containing BPSV40-NLS-HALO from strain YTL1705. To construct plasmid pMvB19 (TEFp-3xHA-TFC4 in pRS315), pRS315 was digested using the SalI and BamHI restriction sites. The *TEF* promoter was amplified from pYM-N18 and 3xHA tagged *TFC4* was amplified from genomic DNA (strain NKI5809). Both PCR products were annealed using an annealing-PCR protocol. The annealed PCR product was cloned into SalI and BamHI digested pRS315 using Gibson Assembly. Subsequently, pMvB20 was constructed by Gibson Assembly of SpeI and NotI digested pMvB19 and an amplified *ADH1* terminator fragment from the pML104 plasmid. To introduce mutations in the *TFC4* gene on pMvB20 (plasmid pMvB21, pMvB22, pMvB23, pMvB24 and pMvB25), pMvB20 was linearized by PCR. Mutant DNA templates were made by amplification of the *TFC4* gene with oligos that harbor point mutations. The linearized plasmid and mutant templates were assembled using Gibson Assembly. For plasmid pMvB24 and pMvB25, two PCR products from *TFC4* were first combined into a single fragment before Gibson Assembly.

### Chromatin immunoprecipitation and qPCR

Chromatin immunoprecipitation (ChIP) was performed as previously described in [30] with the following modifications. Yeast cultures were grown in 75 mL YEPD until mid-log phase (OD_660nm_ = 0.5-0.8). Optionally, cells were switched to repressive conditions by incubating for 2 hours in 2% glycerol (YEPgly) at 30°C or 37°C. For ChIP of Fpt1 and Brf1 in the presence of τ131-mutant plasmids, yeast cells were grown in synthetic complete (SC) media lacking leucine (SC-LEU) to select for the rescue plasmid. For anchor-away, cells were treated with 7.5 µM rapamycin or DMSO for 60 minutes. Cells were cross-linked for 15 minutes with one-tenth of freshly prepared fix solution (11% formaldehyde, 50 mM Hepes-KOH [pH 7.5], 100 mM NaCl, 1 mM EDTA) and cross-linking was quenched for 1 minute with Tris-HCl pH 8.0 (750 mM final concentration). Chromatin was sheared using the Bioruptor PICO (Diagenode) 7 minutes with 30-second intervals at 4°C. For TAP-based IP, 40 µL Dynabeads M-270 Epoxy (ThermoFisher) coupled to rabbit immunoglobulin G (IgG) was mixed with 400 µL chromatin and incubated overnight on a turning wheel at 4°C. For V5-based IP, 1 μL mouse anti-V5 antibody (Invitrogen, R96025) was coupled to 25 μL Protein G Dynabeads (Invitrogen) and incubated overnight on a turning wheel at 4 °C. The next day, the beads were mixed with 400 μL chromatin and again incubated overnight on a turning wheel. Quantitative PCR with technical duplicates was performed as described in [19] using a QuantStudio 5 Real-Time PCR System (Applied Biosystems).

### Live cell imaging

Live cell imaging was performed as previously described [19,31] with minor modifications. Cells were grown to OD_660nm_ 0.2-0.4 in SC + 2% glucose and imaged on a coverslip with an agarose pad consisting of 2% agarose in SC + 2% glucose. Live cell imaging was performed on a setup consisting of an AxioObserver inverted microscope (Zeiss), an alpha Plan-Apochromat 100X NA 1.46 oil objective, an sCMOS ORCA Flash 4v3 (Hamamatsu) with a 475-570 nm dichroic (Chroma), 570 nm longpass beamsplitter (Chroma), 515/30 nm and 600/52 nm emission filters (Semrock), and an UNO Top stage incubator (OKOlab) at 30°C. LED excitation was set at 470/24 nm (for GFP) and 550/15 nm (for mScarlet-I) (SpectraX, Lumencor) at 10% power with an ND1 filter, resulting in a 3.1 W/cm^2^ and 10.3 W/cm^2^, respectively. For each position, a *z*-stack (9 slices, Δ*z* 0.5 µm) was recorded with 200 ms exposure using Micro-Manager software version 1.4 [32]. For each condition, 3-5 images were taken for each of the 3 biological replicates.

### Live cell imaging data analysis

For localization of Fpt1, the microscopy data was analyzed using custom Python software (10.5281/zenodo.7650172). A maximum intensity projection of the mScarlet-I channel was used to segment cells using Otsu thresh-holding and water shedding. For each cell, the total nuclear mScarlet-I intensity was determined in each of the 9 slices in the *z*-stack, and the *z*-slice with the maximum signal was taken to be the slice where the nucleus is in focus. In this *z*-slice, the nuclear and cytoplasmic intensities of the GFP channel were determined and corrected for background by subtracting the median value of all intensities measured in the same z-slice outside the cells. For each cell, the nuclear enrichment was taken to be the ratio between the median nuclear and median cytoplasmic signal.

### Immunoblotting

Immunoblotting and analysis were performed as previously described [19] with minor modifications. Cells were grown in 15 mL YEP or SC-LEU (in case of positive selection for the τ131 plasmids) + 2% glucose or glycerol. The HA-tag was stained with HA primary mouse antibody (Roche 12CA5, 1:2000) and secondary antibody (goat anti-mouse antibody IRDye 800, Licor, 1:10000) for 2h and 45 minutes respectively in 2% milk in TBS-T. Quantification of gels was done using ImageJ software version 1.53g [33] by applying the area under the curve method. Pgk1 protein levels were used to normalize target protein levels.

### Growth assay

To assess growth on solid media, spot test analysis was performed. Yeast cultures were grown overnight in 5 mL SC-LEU to select for plasmid maintenance. Each overnight culture was diluted in SC-LEU to an OD_660nm_ of 1. Serial ten-fold dilutions were subsequently spotted on YEP (with 2% glucose or 2% glycerol) plates containing DMSO or 7.5 µM rapamycin. Growth was assessed after 2 days at 30°C or 37°C to check temperature sensitivity.

### AlphaPulldown and Alphafold multimer predictions

To identify potential protein interaction partners of full-length Fpt1 with RNAPIII and its core transcription machinery, the Alphapulldown pipeline (v1.0.0) was used with default parameters [34]. Representative subunits were selected including RNAPIII components, such as, Rpc1 (P04051), Rpc2 (P22276), Rpc5 (P36121), Rpc9 (P47076), and Rpac1 (P07703); TFIIIB subunits Brf1 (P29056), Bdp1 (P46678); and TFIIIC subunits τ131 (P33339), τ138 (P34111), τ91 (Q06339), τ60 (Q12308), and τ55 (Q12415). We evaluated the Alphafold predicted aligned error (PAE) for residue pairs at interfaces between Fpt1 and its interaction partner using ChimeraX [35] and PAE viewer [36]. Interaction surface regions with the lowest PAE were further analyzed, and residues with the lowest PAE values were selected for mutagenesis in the *TFC4* gene.

### Protein expression and purification of Fpt1Δ234-353

Fpt1 residues 1 to 233 were cloned into the modified pET vector, pETNKI-his-3C-LIC-kan, encoding a 6xhis N-terminal tag followed by a HRV 3C protease recognition site [37]. The construct was transformed into BL21(DE3) cells and the protein was expressed in 1L of LB medium supplemented with 30 µg/mL kanamycin. Cells were grown at 37°C until OD_600nm_ was 0.4 before cells were cooled to 20°C. Expression was induced by addition of 0.4 mM IPTG and cells were grown o/n at 20°C. Cells were harvested by centrifugation for 15 minutes at 4,000 x *g* and the cell pellet was stored at -20°C until further use. For purification, cells were thawed and resuspended into 25 mL lysis buffer (25 mM Tris pH 8.0, 300 mM NaCl, 1 mM TCEP). Cells were lysed by sonication, and the lysate was cleared by centrifugation for 30 minutes at 56,000 x *g* at 4°C. The soluble lysate was collected and loaded onto 2 mL Nickel-chelating beads (Cytiva) in a column by gravity. Beads were washed with 15 mL of wash buffer (25 mM Tris pH 8.0, 300 mM NaCl, 1 mM TCEP, 20 mM imidazole) and protein was eluted in 25 mM Tris pH 8.0, 300 mM NaCl, 1 mM TCEP, 200 mM imidazole. Fractions of 2 mL were collected and analyzed by SDS-PAGE. Fractions containing the protein were pooled and concentrated to 1 mL in a 10 kDa cut-off Vivaspin concentrator. The sample was further purified by size exclusion chromatography using a HiLoad 16/600 S75 column (Cytiva), equilibrated in 25 mM Tris pH 8.0, 150 mM NaCl, 1 mM TCEP, connected to a Bio-Rad NGC system. The protein eluted in a single peak and fractions were pooled and concentrated to 5 mg/mL. Aliquots were flash-frozen in liquid nitrogen and stored at -80°C. The total yield was 14 mg purified Fpt1 core(aa1-233) protein from 1L of bacterial cell culture.

### Protein expression and purification of TFIIICΔtail

The TFIIIC variant lacking the C-terminal region of τ95 that autoinhibits DNA binding activity of τA [10], hereafter referred to as TFIIICΔtail, was generated as previously described [12]. Briefly, the six *S. cerevisiae* TFIIIC subunits, codon-optimized for expression in insect cells, were cloned using the biGBac assembly system [38]. Baculovirus was generated following standard protocols, and the complex was expressed in High Five insect cells using a 1:1000 dilution of the recombinant virus. Cells were harvested at 90–95% viability and stored at −80°C.

For purification, frozen pellets were resuspended in lysis buffer (20 mM HEPES, pH 7.5, 500 mM NaCl, 2 mM MgCl₂, 4 mM β-mercaptoethanol, 10% glycerol). Pellets from 6 liters of culture were used for each purification at a ratio of 3 ml buffer per gram of cell mass. Lysates were stirred on ice, and protease inhibitors (1 tablet per 20 g cells, Sigma–Aldrich), Benzonase (4 μl per 50 ml buffer), and DNase I (500 μl, 10 mg/ml) were added. After full resuspension, the cells were sonicated (3 min, 40% amplitude), and ultracentrifuged at 35,000 rpm for 1 h at 4°C. A volume of 2 ml Strep-Tactin Sepharose™ resin pre-equilibrated in strep-wash buffer (20 mM HEPES, pH 7.5, 150 mM NaCl, 5 mM DTT, 5% glycerol) was added to the supernatant and incubated for 2 h at 4°C. The TFIIICΔtail bound to the beads was washed with 30 ml strep-wash buffer and eluted using strep-elution buffer (strep-wash buffer supplemented with 50 mM biotin).

Eluted TFIIICΔtail was further purified by anion exchange chromatography using a Capto HiRes Q 5/50 column pre-equilibrated with Capto-A buffer (20 mM HEPES, pH 7.5, 150 mM NaCl, 5 mM DTT). Elution was performed using a linear gradient (0–70%) of Capto-B buffer (20 mM HEPES, pH 7.5, 1 M NaCl, 5 mM DTT), followed by a step to 100% Capto-B. Fractions with conductivity values between 25–32 mS/cm, corresponding to TFIIICΔtail, were pooled, buffer exchanged into Capto-A buffer, concentrated, aliquoted, and flash-frozen for storage at −80°C.

### Electrophoretic Mobility Shift Assay

Oligonucleotides (Sigma–Aldrich, HPLC purified) corresponding to both strands of the tRNA^His^ promoter from positions −14 to +76 or DNA mutants were end-labeled using [γ-32P]-adenosine 5′-triphosphate (ATP) and T4 polynucleotide kinase (New England Biolabs) and purified on a 10% acryl/bisacrylamide and 8.3 M (w/v) urea gel. DNA was eluted overnight from excised gel bands in 0.5 M ammonium acetate, 10 mM magnesium acetate, 0.1% (w/v) SDS and 0.1 mM EDTA, and then ethanol precipitated. The labeled strand was annealed with its cold complementary strand at room temperature (RT) for 30 min in 20 mM HEPES (pH 7.5), 5 mM MgCl2 and 100 mM KCl after heat denaturation at 95°C for 3 min. EMSA was performed as follows: DNA (∼30 000 cpm, ∼10 nM) was incubated with increasing amounts of Fpt1(Δ234-353) (0.3 µM to 40 μM) in the presence of 1 μg poly-dI/dC double strand (Amersham) in buffer FB (20 mM HEPES, pH 7.5, 150 mM KCl, 2 mM MgCl_2_, 5 mM DTT) for 1 h at 4°C. For the Fpt1/TFIIICΔtail EMSA, the concentration of TFIIICΔtail was fixed at 0.3 µM and increasing concentration of Fpt1 (Δ234-353) was used (0.3 µM to 30 μM). A control with BSA at the same concentrations was done. After incubation, the samples were loaded onto a native 5% acrylamide gel and run at 4°C at 100 V for 4 h in Tris–glycine buffer. Subsequently, the gel was dried for 1 h at 80°C and the signal visualized by exposure to a phosphorimaging screen.

### Statistical analysis

Statistical details of experiments can be found in the figure legends. In the figure legends, ‘n’ represents the number of biological replicates unless otherwise specified. Statistical analyses were performed using Prism 10 V10.4.1.

## RESULTS

### Fpt1 occupancy at tDNAs depends on both TFIIIC and TFIIIB, independent of nutrient conditions

Fpt1 occupancy at tRNA genes is independent of active RNAPIII transcription but requires both TFIIIB and TFIIIC under transcriptionally active conditions [19]. Notably, the chromatin occupancy levels of Fpt1 resemble those of TFIIIC. Additionally, the dynamic behavior of Fpt1 upon changing nutrient conditions closely mirrors that of TFIIIC, with increased chromatin occupancy under repressive conditions, while TFIIIB occupancy decreases. Strikingly, although Fpt1 shows dynamics opposite to those of TFIIIB, it still requires TFIIIB for chromatin binding under active conditions [19]. To investigate whether Fpt1 remains dependent on both TFIIIC and TFIIIB under repressive conditions, when its occupancy sharply increases, the anchor-away (AA) system was used to conditionally deplete subunits of the essential TFIIIC and TFIIIB complexes from the nucleus [39]. The AA system is based on the rapamycin-induced heterodimerization of human FKBP12 with the FKBP12-rapamycin-binding (FRB) domain of human mTOR [39,40]. To enable this and avoid the toxic stress response of cells to rapamycin, two genetic modifications are required. AA strains lack the *FPR1* gene, encoding a yeast homologue of the human FKBP12 protein, and have a missense mutation in the FRB domain of *TOR1* (*tor1-1*), making Tor1 (and the TORC1 complex in which it resides) resistant to rapamycin treatment but leaving other functions of Tor1 intact. The TORC1 complex is a central regulator of the nutrient response pathway that is inhibited by rapamycin in wild-type cells [41]. TORC1 phosphorylates and activates Sch9, a kinase involved in nutrient signaling downstream of TORC1 [42]. Together with the kinase PKA, Sch9 phosphorylates and thereby inactivates Maf1 [43]. Since TORC1 can be assembled with either Tor1 or its homologue Tor2, which is insensitive to rapamycin, and the *tor1-1* mutation does not affect the TORC1 kinase activity, TORC1 signaling should not be affected in the presence or absence of rapamycin. However, given the central role of TORC1 in RNAPIII regulation via Maf1 and the aim to test Fpt1 dependency on TFIIIB and TFIIIC under repressive conditions, we first validated the stress response to nutrient perturbation in the AA background. We compared the RNAPIII binding dynamics to several tDNA loci in AA reference strains without any anchored protein to the binding dynamics in wild-type non-AA strains. After cells were grown to mid-log phase in glucose-containing media at 30°C, they were either treated with rapamycin (1h) or switched to glycerol-containing media at 37°C, a condition known to repress RNAPIII transcription (**Figure 1A**) [25,44]. For the glycerol condition, cells were treated with rapamycin (1h) after an initial 1h incubation in glycerol at 37°C. As expected, rapamycin treatment of wild-type yeast cells grown in glucose reduced the occupancy of the TAP-tagged largest RNAPIII subunit Rpo31 (% ChIP/input) at selected tRNA genes (**Figure 1B**). In contrast, in the AA strain, which is rapamycin-insensitive, rapamycin treatment did not negatively affect Rpo31 occupancy in glucose compared to DMSO (**Figure 1C**). In both the wild-type and AA background strain, Rpo31 occupancy decreased upon exposure to nutrient stress, and this was not further enhanced by rapamycin treatment. These results confirm that the AA strains display a normal stress response upon nutrient perturbation and that rapamycin does not induce a stress response in these strains.

**Figure 1.**
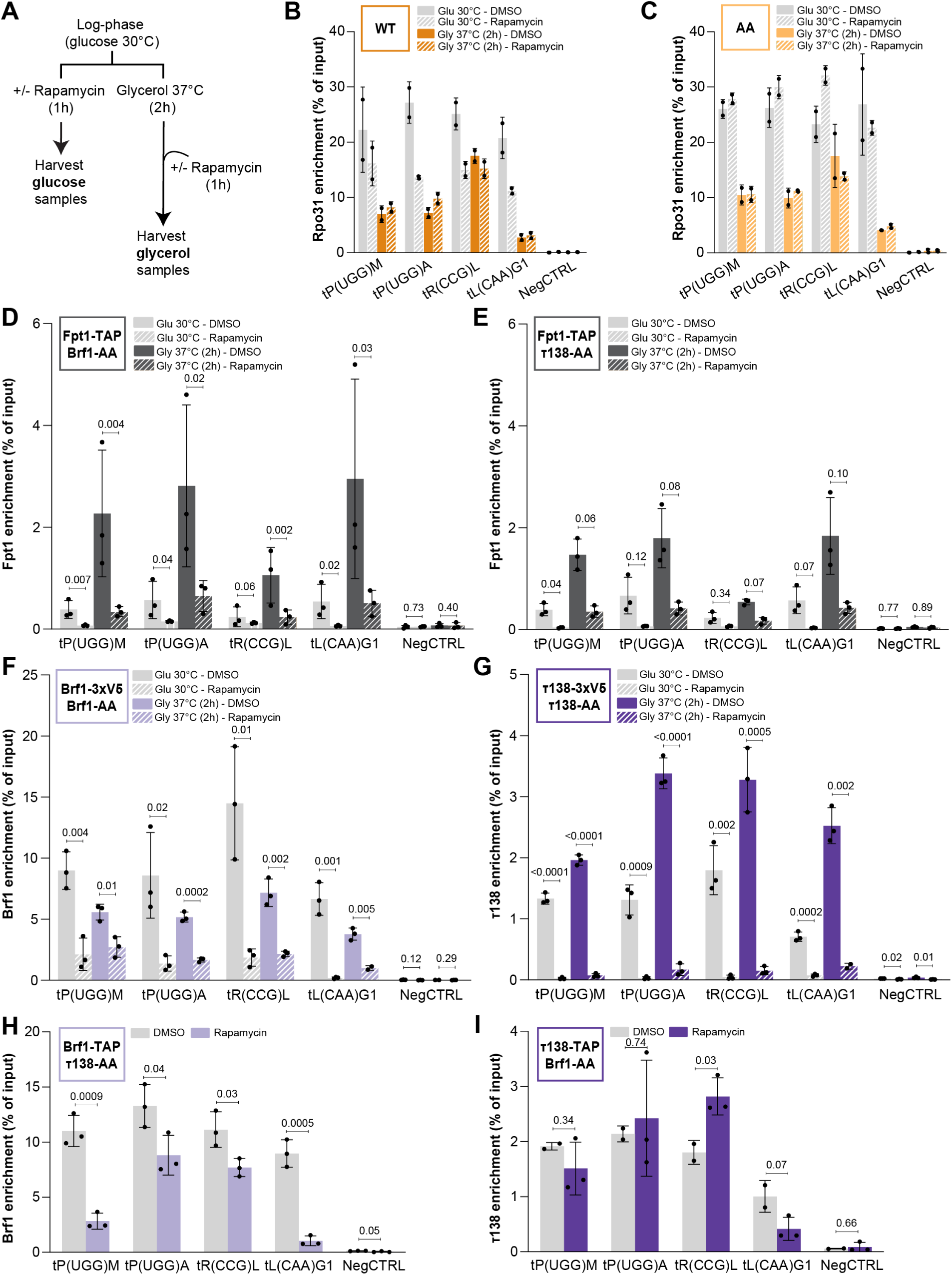
Fpt1 binding at tRNA genes requires both TFIIIB and TFIIIC. a) Outline of experimental set-up: cells were grown to mid-log phase. Glucose samples were immediately treated with 7.5 µM rapamycin or DMSO for 1h. Glycerol samples were first incubated at 37°C for 1h, before addition of rapamycin or DMSO. b) Rpo31 enrichment (TAP-ChIP) in a wild-type background in DMSO and rapamycin-treated cells in both glucose 30°C and 2h glycerol 37°C (n = 2 ± SD). NegCTRL is the negative control locus *ARS504*; an origin of replication devoid of proximal RNAPII and RNAPIII. c) As in (B), Rpo31 TAP-ChIP in an anchor-away (AA) background. d) Fpt1 enrichment (TAP-ChIP) upon nuclear depletion of Brf1 (1h rapamycin) and DMSO control in both glucose 30°C and 2h glycerol 37°C (n = 3 ± SD). e) As in (D), Fpt1 enrichment (TAP-ChIP) upon nuclear depletion of τ138. f) Brf1 enrichment (V5-ChIP) upon nuclear depletion of Brf1 (1h rapamycin) and DMSO control in both glucose 30°C and 2h glycerol 37°C (n = 3 ± SD). g) As in (F), τ138 enrichment (V5-ChIP) upon nuclear depletion of τ138. h) τ138 enrichment (TAP-ChIP) upon nuclear depletion of Brf1 (1h rapamycin) and DMSO control in glucose 30°C (n = 3 ± SD). i) As in (H), Brf1 enrichment (TAP-ChIP) upon nuclear depletion of τ138.

Next, we employed the anchor-away system to conditionally deplete TFIIIB (Brf1-AA) and TFIIIC (τ138-AA) in conditions of both active and inactive RNAPIII transcription. In conditions of active RNAPIII transcription (glucose 30°C), nuclear depletion of Brf1 and τ138 resulted in a loss of Fpt1 binding (**Figure 1D-E**), consistent with our previously published findings [19]. Under repressive conditions, which cause an increase in Fpt1 occupancy [19], nuclear depletion of Brf1 and τ138 diminished this increase in Fpt1 occupancy (**Figure 1D-E**). Interestingly, after depletion of TFIIIB and TFIIIC under repressive conditions, Fpt1 occupancy levels were comparable to those observed in glucose (DMSO). Previous studies have shown that total Fpt1 protein levels increase under repressive conditions [19], which could lead to higher association with tDNAs. However, increased protein levels using a constitutive promoter did not increase tDNA binding, suggesting other mechanisms must be at play [19]. A second possibility is that nuclear depletion of TFIIIB and TFIIIC is less efficient in glycerol at 37°C compared to glucose at 30°C. Alternatively, Fpt1 has intrinsic, TFIIIB- and TFIIIC-independent tDNA binding capabilities under repressive conditions, but tDNA binding is more efficient in the presence of TFIIIB and TFIIIC. To test the second scenario, the depletion efficiency of Brf1 and τ138 was assessed under both active and repressive conditions (**Figure 1F-G**). Occupancy of Brf1 and τ138 was reduced in both glucose at 30°C and glycerol at 37°C. However, Brf1 was depleted from chromatin less efficiently than τ138, possibly due to the longer residency time of TFIIIB on chromatin [45] or limited accessibility of the FRB-tag within the folded TFIIIB complex. Since Fpt1 binding was higher in glycerol than glucose while Brf1 binding was similar upon depletion in both conditions, these results suggest that Fpt1-tDNA interactions are partially independent of TFIIIB in repressive conditions.

Depletion of τ138 and Brf1 showed similar effects on Fpt1 binding (**Figure 1D-E**), suggesting that Fpt1 relies on both TFIIIB and TFIIIC. To investigate whether TFIIIC depletion affects Fpt1 indirectly through TFIIIB depletion and vice versa, we examined the interdependency of TFIIIC and TFIIIB under the conditions analyzed (**Figure 1H-I**). Upon nuclear depletion of Brf1, τ138 remained capable of binding to the tRNA genes analyzed, while Fpt1 binding was abolished (**Figure 1D and H**). In contrast, upon nuclear depletion of τ138, Brf1 occupancy was reduced but not abolished at RNAPIII-transcribed genes (**Figure 1E and I**). Of note, the extent of Brf1 reduction upon τ138 depletion varied among individual tRNA genes with some genes showing only minor changes in Brf1 binding. Taken together, these results suggest that optimal Fpt1 binding to tDNAs requires both TFIIIC and TFIIIB, independent of each other, and independent of nutrient availability.

### The C-terminal region of Fpt1 is crucial for nuclear localization and increased tDNA occupancy under repressive conditions

Given the requirement of both TFIIIB and TFIIIC for increased binding of Fpt1 under repressive conditions, we further investigated how different domains in Fpt1 contribute to its interaction with tDNAs. To identify functional regions in Fpt1 that are essential for stress-induced tDNA binding, several truncation mutants were generated based on the predicted protein structure from AlphaFold (AlphaFold Monomer v2.0 pipeline, Uniprot: Q02209) [46,47] (**Figure 2A-B, Supplemental Figure 2A**). Based on the AlphaFold model confidence scores (pLDDT) and recent definitions of protein disorder [48], the following domains were defined: a structured core domain (residue 1-233), and an intrinsically disordered region (IDR, residue 234-353) that contains a potentially structured α-helix (residue 290-314) (**Figure 2A**). In addition, based on several NLS-prediction programs, a potential nuclear localization signal (NLS) was found [49–51] located near the C-terminus.

**Figure 2.**
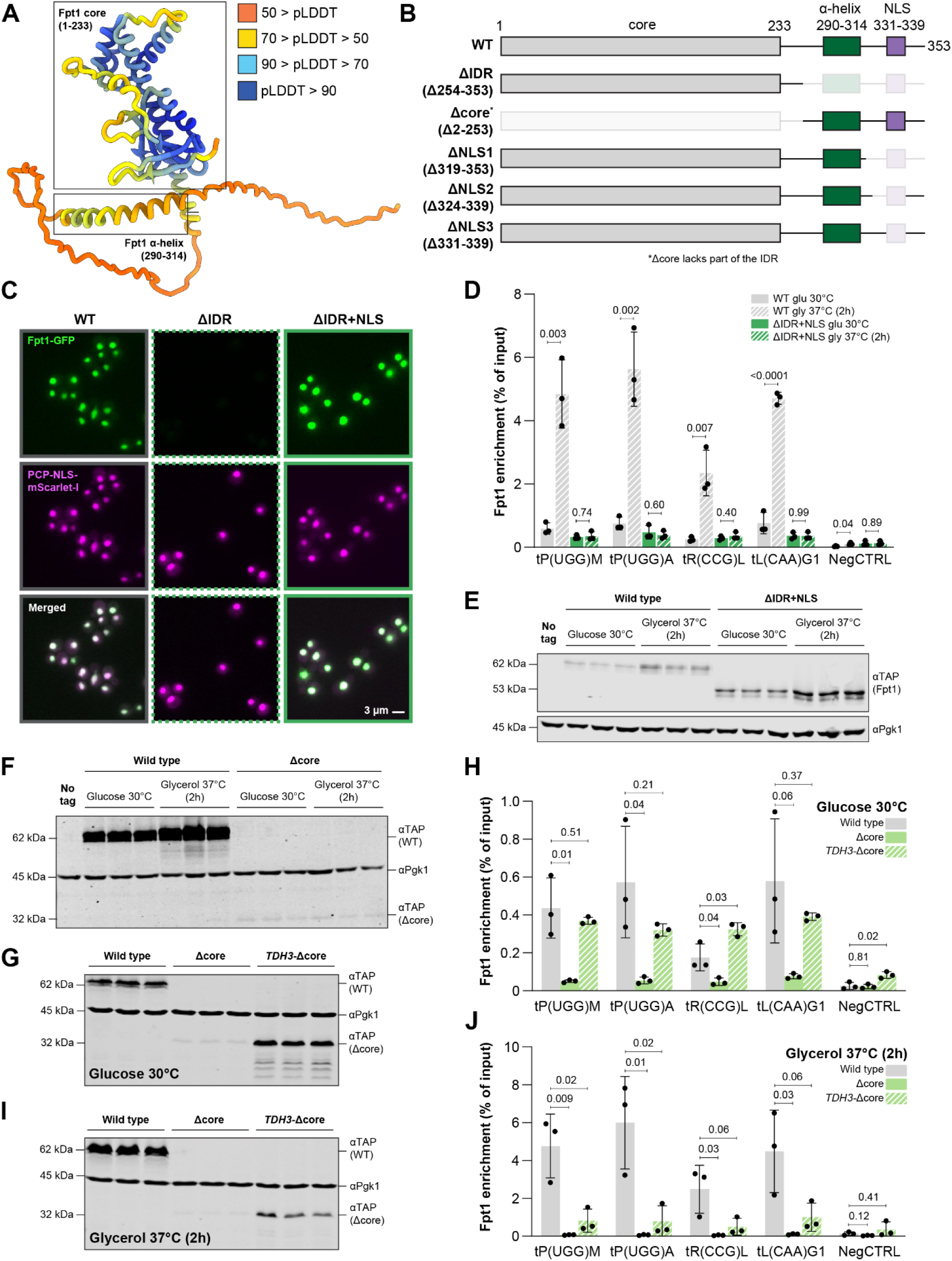
The IDR of Fpt1 encodes a nuclear localization signal and stress-responsive element, while the core domain ensures protein stability. a) AlphaFold2-predicted protein structure of Fpt1 (AlphaFold Monomer v2.0 pipeline, Uniprot: Q02209). The confidence of the pLDDT (predicted local distance difference test) is shown by different colors. b) Schematic representation of several truncation mutants of Fpt1. IDR = intrinsically disordered region, NLS = nuclear localization signal. c) Representative images of wild-type Fpt1, ΔIDR, and ΔIDR+NLS localization in live yeast cells in glucose 30°C. From top to bottom: Fpt1-GFP (green), PCP-NLS-mScarlet-I as a nuclear marker (magenta), and the merged channel. Scale bar: 3 μm. d) Fpt1 enrichment (TAP-ChIP) of wild type and ΔIDR+NLS in glucose 30°C and 2h glycerol 37°C (n = 3 ± SD). e) Fpt1-TAP immunoblot (n = 3) of wild type and ΔIDR+NLS in glucose 30°C and 2h glycerol 37°C. No-tag = BY4741. f) Fpt1-TAP immunoblot (n = 3) of wild type and Δcore in glucose 30°C and 2h glycerol 37°C. g) Fpt1-TAP immunoblot (n = 3) of wild type, Δcore, and *TDH3-*Δcore in glucose 30°C. h) Fpt1 enrichment (TAP-ChIP) of wild type, Δcore, and *TDH3-*Δcore in glucose 30°C (n = 3 ± SD). i) As in (G), in 2h glycerol 37°C. j) As in (H), in 2h glycerol 37°C.

First, the role of the C-terminal IDR was examined. Since wild-type Fpt1 is a nuclear protein, the cellular localization of a truncation mutant that lacked most of the intrinsically disordered C-terminus (Δ254-353, termed Fpt1ΔIDR) was tested using fluorescence microscopy of endogenously GFP-tagged versions of Fpt1 (**Figure 2C, Supplemental Figure 2B**). In line with the predictions that an NLS is located in the C-terminal IDR, Fpt1ΔIDR was defective in nuclear localization. By testing multiple truncation lengths, the region essential for the NLS function was mapped to residues 331-339 (**Supplemental Figure 2C-D**). To study the nuclear function of the Fpt1ΔIDR mutant, a bipartite SV40 NLS was attached to the C-terminus to restore nuclear localization in the absence of the native NLS [52]. Fpt1ΔIDR+NLS was localized in the nucleus and remained bound to tRNA genes above background levels under conditions of active transcription. However, Fpt1ΔIDR+NLS failed to increase tDNA occupancy under repressive conditions (2h glycerol 37°C) (**Figure 2C-D**). This observation could not be explained by lower protein levels, as Fpt1ΔIDR+NLS was expressed at higher levels compared to wild type in both active (4.6-fold) and repressive (2.9-fold) conditions (**Figure 2E, Supplemental Figure 2E**). Therefore, these results suggest that the C-terminal IDR is essential for stress-induced enrichment of Fpt1 at tRNA genes and for Fpt1 destabilization.

To test whether the core domain also contributes to increased Fpt1 binding upon nutrient stress, a truncation mutant lacking the core domain, including a small part of the IDR (Δ1-253), termed Fpt1Δcore, was constructed. As expected, since the NLS was mapped to residue 331-339, Fpt1Δcore was localized in the nucleus (**Supplemental Figure 2F**). However, TAP-tagged Fpt1Δcore was poorly expressed (**Figure 2F, Supplemental Figure 2G**). In an attempt to restore Fpt1Δcore protein levels to wild type and to be able to assess its dynamics at tRNA genes, a strong constitutive *TDH3* promoter was inserted upstream of the *FPT1* gene. The *TDH3* promoter was able to restore Fpt1Δcore protein levels in glucose at 30°C (**Figure 2G, Supplemental Figure 2H**), resulting in tDNA binding comparable to that of wild type (**Figure 2H**). However, *TDH3* promoter-driven Fpt1Δcore did not restore binding in glycerol at 37°C and was only partially able to increase Fpt1Δcore protein levels under this condition (**Figure 2I-J, Supplemental Figure 2I**). The lower protein levels might at least in part explain the reduced binding of Fpt1 without its core domain. Taken together, we established that the core domain is important for protein stability and contributes to basal Fpt1 recruitment in glucose, while the IDR is essential for increased Fpt1 occupancy at tRNA genes under repressive conditions.

### A short predicted α-helix in the C-terminal region of Fpt1 is required for Fpt1 recruitment to tRNA genes

Fpt1 requires both TFIIIB and TFIIIC to interact with tRNA genes and the C-terminal IDR in Fpt1 is crucial for increased Fpt1 binding during nutrient stress. These findings raise the possibility that Fpt1 occupancy increases under repressive conditions through interactions between the IDR of Fpt1 and components of TFIIIB or TFIIIC. To screen for candidate protein-protein interactions between Fpt1 and core components of the RNAPIII transcription machinery, AlphaPulldown was used [34]. The highest-confidence interaction was predicted between Fpt1 and τ131, a subunit of the τA subcomplex of TFIIIC (**Figure 3A, Supplemental Figure 3A**). A predicted structured C-terminal α-helix within the IDR of Fpt1 was predicted to interact with the conserved N-terminal TPR array in τ131. This region in τ131 is known to interact with Brf1 to facilitate assembly of TFIIIB at tRNA genes [16,53,54]. Notably, the C-terminal IDR of Fpt1 increased tDNA occupancy under repressive conditions (**Figure 2D**) and Fpt1’s binding efficiency and dynamics resemble those of TFIIIC. We therefore hypothesized that the C-terminal α-helix mediates Fpt1’s interaction with τ131, establishing its dependence on TFIIIC and consequently contributing to increased tDNA binding under repressive conditions. To test the importance of the C-terminal α-helix, an Fpt1 mutant lacking the structured C-terminal α-helix (Δ290-314) termed Fpt1Δα-helix (**Figure 3B**) was made. Removal of the α-helix did neither affect nuclear localization nor protein levels (**Figure 3C-E**). Similar to the Fpt1ΔIDR mutant, deletion of the α-helix abolished the increased binding of Fpt1 under repressive conditions (2h glycerol 37°C) (**Figure 3F**). Therefore, the structured C-terminal α-helix could be considered the stress responsive element in Fpt1, hereafter named the SRα (Stress Responsive α-helix). Interestingly, deletion of α-helix also decreased binding of Fpt1 in glucose (**Figure 3F**), suggesting that apart from being the stress-responsive element, the SRαx may also have a role in Fpt1 binding under active conditions.

**Figure 3.**
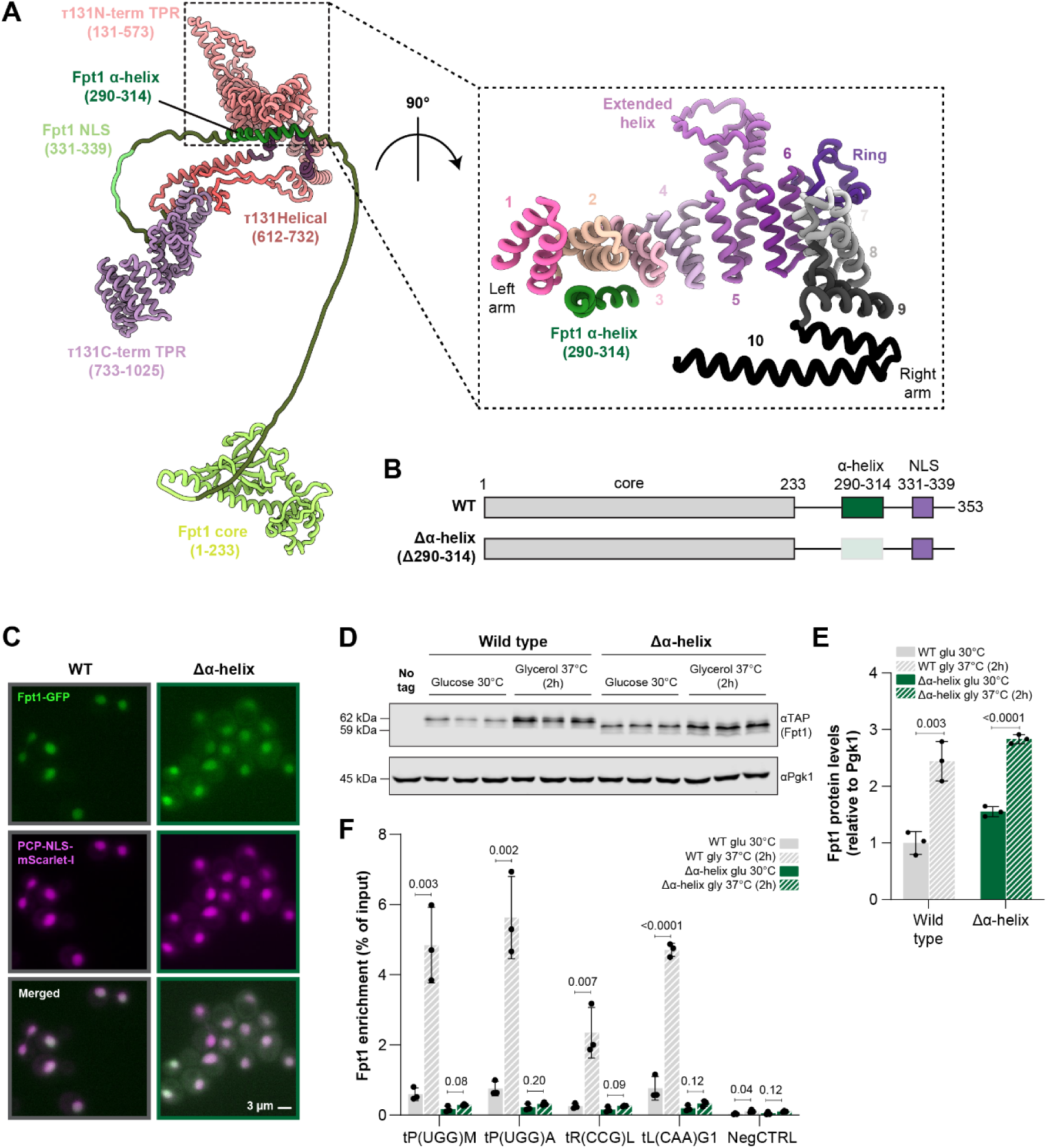
A small region in the disordered region is predicted to interact with τ131 and responsible for the stress response. a) Left panel: AlphaFold2-predicted interaction between Fpt1 and TFIIIC-subunit τ131. Fpt1 is colored according to its domain organization: Fpt1 core (residues 1–233), Fpt1 α-helix (290–314), and Fpt1 NLS (331–339). The interacting partner, τ131, is divided into three regions: the N-terminal TPR domain (residues 131–573), the helical region (612–732), and the C-terminal TPR domain (733–1025). Right panel: A close-up of the interface between the Fpt1 α-helix and the τ131 N-terminal TPR, corresponding to the boxed region. TPR repeat numbers and right and left arm are indicated. b) Schematic representation of wild-type Fpt1 and the Δα-helix mutant. c) Representative images of wild-type Fpt1 and Δα-helix localization in live yeast cells in glucose 30°C. From top to bottom: Fpt1-GFP (green), PCP-NLS-mScarlet-I as a nuclear marker (magenta), and the merged channel. Scale bar: 3 μm. d) Fpt1-TAP immunoblot (n = 3) of wild type and Δα-helix in glucose 30°C and 2h glycerol 37°C. No-tag = BY4741. e) Quantification of (D). f) Fpt1 enrichment (TAP-ChIP) of wild type and Δα-helix in glucose 30°C and 2h glycerol 37°C (n = 3 ± SD).

### Increased tDNA occupancy of Fpt1 in stress conditions is mediated through interactions with τ131

The SRα in Fpt1 is required for its dynamic response upon nutrient perturbation. In addition, this SRα is predicted to interact with the TFIIIC subunit τ131. Notably, TFIIIC occupancy at tRNA genes increases under repressive conditions, similar to Fpt1. Based on these findings, we hypothesized that the interaction between the N-terminal TPR array in τ131 and the SRα in Fpt1 facilitates increased Fpt1 binding under repressive conditions. To test this hypothesis, we aimed to mutate τ131 to disrupt its putative interaction with the SRα in Fpt1. However, the N-terminal TPR array in τ131 is a conserved domain important for TFIIIB recruitment and mutations in this region could potentially affect TFIIIB recruitment and cell viability [55,56]. To avoid these issues, a strategy based on the AA system in combination with wild-type and mutant rescue plasmids was used (**Figure 4A**). Endogenous τ131 was fused with the AA FRB-tag and conditionally depleted using the AA system while a rescue plasmid harboring a HA-tagged wild-type copy of τ131 was introduced to restore the loss of the endogenous copy. This approach relies on the use of protein-tags to deplete the endogenous τ131 copy (FRB-tag), and detect the plasmid τ131 copy (HA-tag). However, in our previous Epi-Decoder dataset, C-terminally TAP-tagged τ131 was not found enriched at the barcoded tRNA gene [19]. Epitope tags can negatively affect protein function [57] and the C-terminus of τ131 is buried within the τA complex, thereby potentially hindering the accessibility of a C-terminal tag [10,12]. Therefore, we created an N-terminally TAP-tagged allele of τ131, which showed a higher expression level and occupancy at tRNA genes than C-terminally TAP-tagged τ131 (**Figure 4B-C**). Subsequent experiments were therefore performed with N-terminally HA-tagged τ131 expressed from a plasmid and an N-terminal FRB-tag for depletion of the endogenous copy.

**Figure 4.**
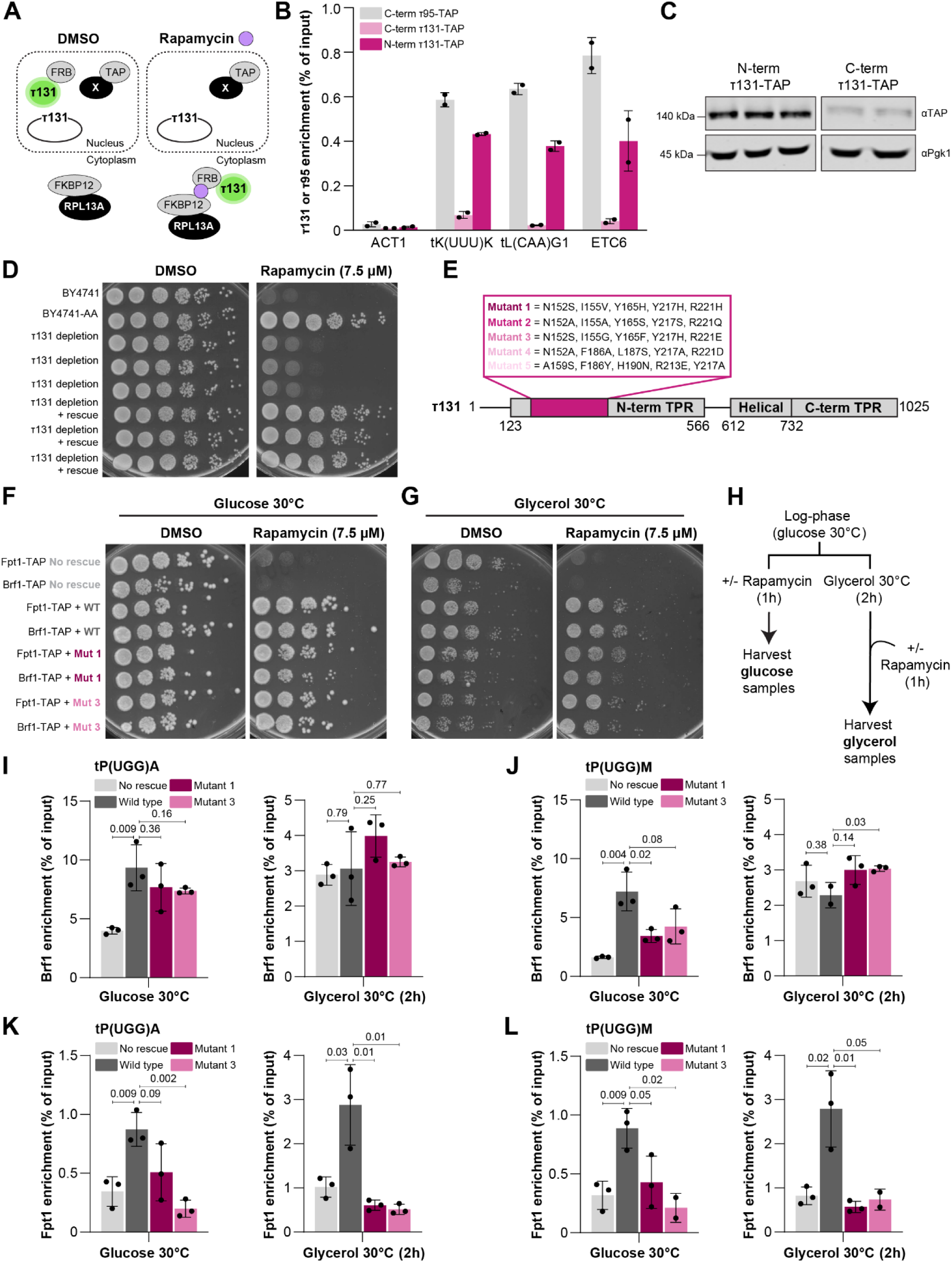
The N-terminal TPR array of τ131 mediates the stress response of Fpt1. a) Schematic representation of the anchor away (AA) and τ131 rescue approach. Endogenous GFP-FRB-tagged τ131 is depleted from the nucleus after 1h rapamycin (7.5 μM) treatment. Loss of endogenous τ131 is complemented with a plasmid-based copy of τ131. X-TAP refers to either Brf1-TAP or Fpt1-TAP. b) N- and C-terminally TAP-tagged τ131 and τ95 enrichment (TAP-ChIP) in glucose 30°C (n = 2 ± SD) at the two tRNA genes *tK(UUU)K* and *tL(CAA)G1* and ETC-site *ETC6*. Τ95 functions as a reference for tDNA binding. *ACT1* is a negative, RNAPII-transcribed locus. c) Immunoblot of N-terminally TAP-tagged τ131 (n = 3) and C-terminally TAP-tagged τ131 (n = 2). d) Spot test analysis on rapamycin- (7.5 µM) and DMSO-containing YEP + 2% glucose plates at 30°C. τ131 depletion is endogenous τ131 depletion without rescue. τ131 depletion + rescue is endogenous τ131 depletion in the presence of a τ131 rescue plasmid. e) Schematic representation of the five τ131 mutants tested. f) Spot test analysis on rapamycin- (7.5 µM) and DMSO-containing YEP + 2% glucose plates at 30°C. No rescue = empty vector without a τ131 copy, WT = wild type τ131 plasmid copy, Mut 1 = τ131 mutant 1, Mut 3 = τ131 mutant 3. g) As in (F), on YEP + 2% glycerol plates at 30°C. h) Outline of experimental set-up: cells were grown to mid-log phase. Glucose samples were immediately treated with 7.5 µM rapamycin for 1h. Glycerol samples were first incubated at 30°C for 1h, before addition of rapamycin or DMSO. i) Brf1 enrichment (TAP-ChIP) in a no rescue (empty plasmid), wild type (wild type τ131 plasmid copy), mutant 1 and mutant 3 (mutated τ131 plasmid copy) background at *tP(UGG)A* upon rapamycin treatment in glucose 30°C (left) and 2h glycerol 30°C (right). n = 3 ± SD. j) As in (I), Brf1 enrichment (TAP-ChIP) at *tP(UGG)M*. k) As in (I), Fpt1 enrichment (TAP-ChIP). l) As in (J), Fpt1 enrichment (TAP-ChIP).

As a proof of principle, we first confirmed that depletion of endogenous τ131 could be rescued by a stable single-copy plasmid expressing the wild-type *TFC4* gene (encoding τ131) from the constitutive *TEF1* promoter. A growth assay on rich media containing glucose showed that continuous τ131-AA depletion without rescue caused lethality when grown on rapamycin-containing plates (**Figure 4D**). A rescue plasmid harboring wild-type *TFC4* restored cell viability similar to the BY4741-AA control strain suggesting successful rescue of endogenous τ131 depletion. To disrupt the predicted interaction between the SRα domain in Fpt1 and the TPR array in τ131, five mutant constructs were designed, each harboring five amino acid substitutions, varying in their predicted potential to disrupt the Fpt1-τ131 interaction (**Figure 4E**). The mutants were designed in such a way that they would disrupt the predicted interaction with Fpt1 and minimally affect TFIIIB recruitment. To identify mutants that fulfill the requirement of minimal TFIIIB disruption, an initial n = 1 screen for Brf1 binding in both active (glucose 30°C) and repressive (2h glycerol 37°C) conditions was performed (**Supplemental Table S2**). As controls, a plasmid without a *TFC4* copy (no rescue) and a plasmid with a wild-type *TFC4* copy (wild type) were taken along. For tRNA genes *tP(UGG)A* and *tP(UGG)M*, τ131 mutant 3 had the smallest effect on Brf1 binding under active conditions when compared to the wild-type copy, and mutant 1 performed second to best (**Supplemental Table S2**). Since τ131 mutants 1 and 3 showed the smallest effect on TFIIIB binding, they were used in subsequent experiments. These mutants restored viability to comparable levels as the wild type rescue upon continuous depletion of endogenous τ131 under active conditions (glucose at 30°C), further supporting the notion that these τ131 mutants are proficient in interacting with TFIIIB (**Figure 4F**).

Having identified two candidate τ131 mutants with a modest effect on TFIIIB binding and viability, we validated these findings and investigated if the mutations block the recruitment of Fpt1 under active and repressive conditions using independent replicates. A commonly used condition to induce repressive stress conditions is growth in glycerol at 37°C. However, the fact that the Brf1 binding levels in glycerol at 37°C were similar across all mutants and comparable to those of the empty vector (**Supplemental Table S2**) led us to ask whether the mutants might be temperature sensitive. We noted that the τ131 mutants showed decreased viability at 37°C (**Supplemental Figure 4A**). Of note, Brf1-TAP appeared to function as a hypomorphic allele, as its viability in combination with τ131 mutants was more reduced compared to that of Fpt1-TAP. The temperature sensitivity of the τ131 mutants is not unprecedented, as multiple temperature sensitive TFIIIC mutants have previously been described [10,58–60]. Therefore, rescue experiments under repressive conditions were executed in glycerol at 30°C. A growth assay confirmed that in glycerol at 30°C, the τ131 mutants did not have detectable growth defects compared to the wild-type rescue (**Figure 4G**). Finally, to test whether the τ131 protein levels were affected by the mutations in the N-terminal TPR-array, immunoblotting was performed (**Supplemental Figure 4B**). Expression of the τ131 rescue copy under the control of a constitutive *TEF* promoter was higher than endogenously tagged τ131 but expression of the mutant forms of τ131 was not different from the wild-type plasmid copy.

To assess Fpt1 and Brf1 binding, cells were grown in glucose-containing media at 30°C until mid-log phase and subsequently treated with rapamycin for 1 h or switched to glycerol-containing media at 30°C. After 1 h in glycerol, cells were treated with rapamycin for 1 h (**Figure 4H**). The τ131 mutants 1 and 3 only mildly affected Brf1 occupancy at the *tP(UGG)A* gene under active conditions (glucose 30°C) (**Figure 4I**). While Brf1 binding was mildly affected at *tP(UGG)A*, the τ131 mutants caused a decrease in Fpt1 binding in glucose 30°C (**Figure 4K**). Interestingly, for the *tP(UGG)M* gene, the effect on Brf1 occupancy under active conditions was larger (**Figure 4J**). Therefore, for this gene we cannot exclude that the effect of the τ131 mutants on Fpt1 binding was mediated in part by loss of Brf1 binding (**Figure 4L**). Under repressive conditions, the τ131 mutants and plasmid without a *TFC4* copy (no rescue) did not affect Brf1 binding, suggesting Brf1 is able to bind independently of τ131 under these conditions in this time interval (**Figure 4I-J**). While TFIIIB binding was not consistently affected by the τ131 mutants under repressive conditions (nor were its protein levels, **Supplemental Figure 4C-D**), Fpt1 binding was significantly reduced (**Figure 4K-L**) even though its protein levels were minimally affected (**Supplemental Figure 4E-F**), further uncoupling the binding dynamics of Fpt1 from that of Brf1.

Taken together, mutations in the N-terminal TPR array of τ131 impaired Fpt1 occupancy at tRNA genes under both active and repressive conditions. Under active conditions, TFIIIB binding was largely unaffected in the τ131 mutants, suggesting that the mutations disrupted Fpt1 recruitment at selected tRNA genes without compromising TFIIIB assembly to the same extent. Under repressive conditions, TFIIIB binding appeared to be independent of the τ131 mutants, yet Fpt1 binding was significantly reduced. These results indicate that the mutated residues in τ131 contribute specifically to the increased occupancy of Fpt1 in response to nutrient perturbation.

## DISCUSSION

Here we combined protein depletion, structural predictions, and mutational studies to understand the interactions between Fpt1 and the core RNAPIII transcription machinery and how these interactions contribute to its recruitment to tRNA genes. Our results demonstrate that Fpt1 requires both TFIIIB and TFIIIC to bind tRNA genes, independent of each other and of nutrient availability. Various mutants of Fpt1 showed that the core domain (aa1-253) and IDR (aa254-353) contribute differently to protein stability and chromatin occupancy. While the core domain stabilizes the Fpt1 protein and is able to interact with tDNAs at basal levels under active conditions, the C-terminal region is important for increased tDNA occupancy under repressive conditions. *In silico* pulldown of Fpt1 with core components of the RNAPIII transcription machinery pointed to a putative interaction between the N-terminal TPR array of τ131 and a structured α-helix embedded within the C-terminal IDR of Fpt1. Similar to deletion of the complete IDR in Fpt1, a mutant lacking the α-helix (named SRα, Stress Responsive α-helix) was unable to increase chromatin occupancy under repressive conditions. Subsequent studies of τ131 mutants, harboring amino acid substitutions in the predicted Fpt1-interaction region, impaired increased binding of Fpt1 under repressive conditions. These results support a model in which the SRα in Fpt1 interacts with τ131 to facilitate stress-induced recruitment of Fpt1 to tDNAs. Interestingly, deletion of the SRα as well as the τ131 mutants also reduced Fpt1 occupancy at tDNAs under active conditions. This suggests that the SRα is not only required for stress-induced Fpt1 recruitment, but also contributes to basal chromatin association, possibly through interactions with TFIIIC. Alternatively, the SRα could stabilize the overall Fpt1 protein conformation (without affecting protein expression), promoting its chromatin-binding capabilities.

### A dual-mode model for Fpt1 recruitment to tRNA genes

Based on our results, Fpt1 recruitment to tRNA genes likely occurs through multiple mechanisms, dependent on nutrient availability (**Figure 5**). Under active conditions, Fpt1 binds tDNAs through its structured core domain, which may interact with TFIIIB or TFIIIC. In support of the latter, using a purified truncated protein of Fpt1 (Fpt1Δ234-353, we could not obtain purified full length Fpt1 protein), *in vitro* electrophoretic shift mobility assay (EMSA) showed a shift in the TFIIIC-tDNA complex upon addition of Fpt1Δ234-353 (**Supplemental Figure 4G**). This suggest that the core domain of Fpt1 is able to interact with at least the TFIIIC complex. Under repressive conditions, TFIIIC occupancy at tRNA genes rapidly increases while TFIIIB occupancy decreases, especially after prolonged periods of stress [22]. Simultaneously, Fpt1 occupancy increases, likely through interactions between its SRα domain in the C-terminal tail with the TFIIIC subunit τ131. This model emphasizes two distinct domains within Fpt1 that mediate context-specific interactions with the RNAPIII core machinery. On the one hand, the structured core establishes basal chromatin interactions through TFIIIB, TFIIIC or direct interaction with the DNA; on the other hand, the SRα facilitates increased chromatin occupancy under repressive conditions through interactions with TFIIIC.

**Figure 5.**
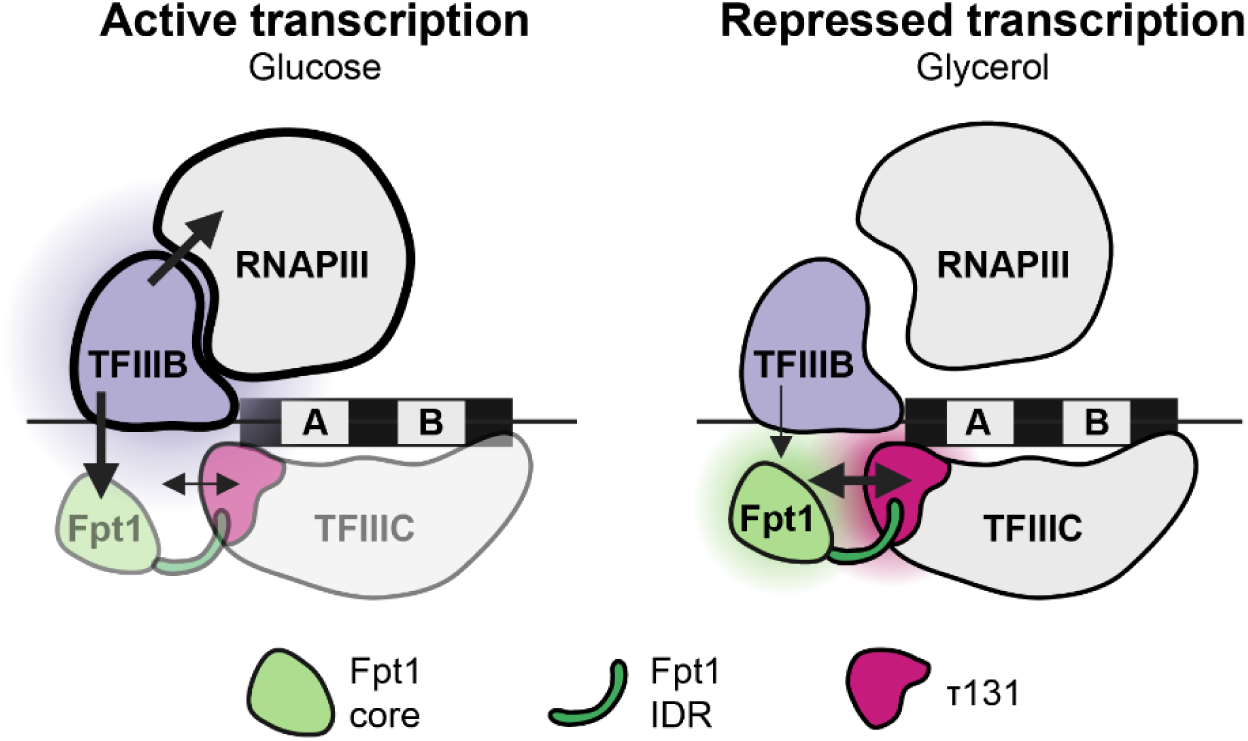
A dual-mode model for Fpt1 recruitment to tRNA genes. In conditions of active transcription (glucose), tRNA genes are pre-dominantly occupied by TFIIIB and RNAPIII. Fpt1 binds tDNAs through its structured core domain, and this binding depends on TFIIIB and TFIIIC. Upon a switch to conditions of repressed transcription (glycerol), TFIIIC occupancy increases while TFIIIB and RNAPIII occupancy decreases. Simultaneously, Fpt1 occupancy increases, likely through interactions between its SRα domain within the C-terminal tail with TFIIIC-subunit τ131. Created with BioRender.com.

### TFIIIB binding in TFIIIC mutants

The N-terminal TPR array in τ131 contains 10 TPRs that are separated into two arms (**Figure 3A**) [15]. Previously described mutations in τ131 that increase RNAPIII transcription [55] have been primarily mapped to TPR2 on the left arm, while mutations that decrease RNAPIII transcription [54] are mapped to TPRs 8-10 on the right arm. Several *in vitro* studies have shown that mutations in TPR2 enhance Brf1 binding and the N-terminus (Nt) of τ131 is required for high-affinity Brf1 binding [15,55], which suggest that the binding site for Brf1 likely exists within Nt-TPR5. Here, the tested τ131 mutants, that each have five amino acid substitutions in TPR1-3, did not show increased binding of Brf1 at the tRNA genes *tP(UGG)M* and *tP(UGG)A* (**Figure 4I-J**). However, unlike *in vitro* studies that examined the effect of single amino acid substitutions [15], we introduced five substitutions simultaneously. The absence of increased Brf1 binding in our experiments may therefore result from the additive effects of these five substitutions.

### New insights into dynamics of the core RNAPIII transcription machinery

The notion that tRNA genes are a homogenous group of genes has been challenged in yeast and humans [61–65]. In line with this, our anchor-away experiments showed that the degree to which sustained TFIIIB binding depends on TFIIIC varies between different tDNAs (**Figure 1H-I**). Indeed, the τ131 mutants also showed tDNA-specific effects on TFIIIB binding under active conditions (**Figure 4I-J**). These tDNA-specific effects suggest that the dynamics of the core RNAPIII transcription machinery are context-dependent and supports the hypothesis that tRNA genes are non-homogenously regulated. Especially interesting is the observation of different TFIIIB and TFIIIC dynamics at tRNA genes *tP(UGG)*M and *tP(UGG)A* (**Figure 1H-I**, **Figure 4I-J**). These tRNA genes have the exact same body sequence and hence the same internal promoter sequence, but have different flanking regions. This suggests that gene position and flanking sequences play a role in the differential dynamics of TFIIIB, TFIIIC, but possibly also of Fpt1. Interestingly, under repressive conditions, TFIIIB binding remained largely unaffected in τ131 mutants or upon depletion of τ131 during the time scale of the experiment (**Figure 4I-J**). Perhaps, under these conditions, TFIIIB adopts a different chromatin-bound state that is less dependent on TFIIIC. This is in line with *in vitro* evidence suggesting that once TFIIIB is assembled at tDNAs, it remains stably bound independent of TFIIIC [13]. Alternatively, the TFIIIB enrichment detected under repressive conditions may reflect non-functional residual binding or crosslinking artifacts. Nonetheless, these results suggest a decoupling of TFIIIB binding from TFIIIC upon prolonged nutrient stress.

## DATA AVAILABILITY

Original western blot images and microscopy data have been deposited at Mendeley Data (DOI: 10.17632/9g7jd5jxc6.1) and are publicly available as of the date of publication. Original code to analyze microscopy data has been deposited at Zenodo (DOI: https://doi.org/10.5281/zenodo.7650171) and is publicly available as of the date of publication. All processed data are in the main text or the supplementary materials.

## ACKNOWLEDGEMENTS

This work was supported by an institutional grant of the Dutch Cancer Society and of the Dutch Ministry of Health, Welfare and Sport, by ZonMW (grant ZonMW open 09120232310068 to FvL), by Oncode Institute (TLL), which is partly financed by the Dutch Cancer Society, and by the Dutch Research Council (VIDI: VI.Vidi.213.031 to TLL). WS-D acknowledges support from the EMBL International PhD program. WS-D, FB and CWM acknowledge support by EMBL. The funders had no role in study design, data collection and interpretation, or the decision to submit the work for publication. We thank Jonas Weidenhausen for advice on the τ131 mutations. We thank Alex Fish for discussions about Fpt1 *in vitro* protein experiments. We thank the Research High Performance Computing Facility of the NKI for assistance. We thank members of the FvL and CWM labs for helpful discussions. We acknowledge the following resources: Yeast Genome Database [66] and BioRender.com.

## AUTHOR CONTRIBUTIONS

Conceptualization: MEvB, FvL

Funding acquisition: FvL, TLL, CWM

Investigation and Methodology: MEvB, WS-D, MJB, JVWM, IAH, MNC, TvW, DA, FB, PC

Supervision: FvL, TLL, CWM

Writing: MEvB, WS-D, MJB, CWM, FvL

