## Supplementary Figures for "Recruitment of Fpt1 to tRNA genes requires TFIIIB and the N-terminal TPR array of TFIIIC subunit τ131"

SUPPLEMENTAL FIGURES

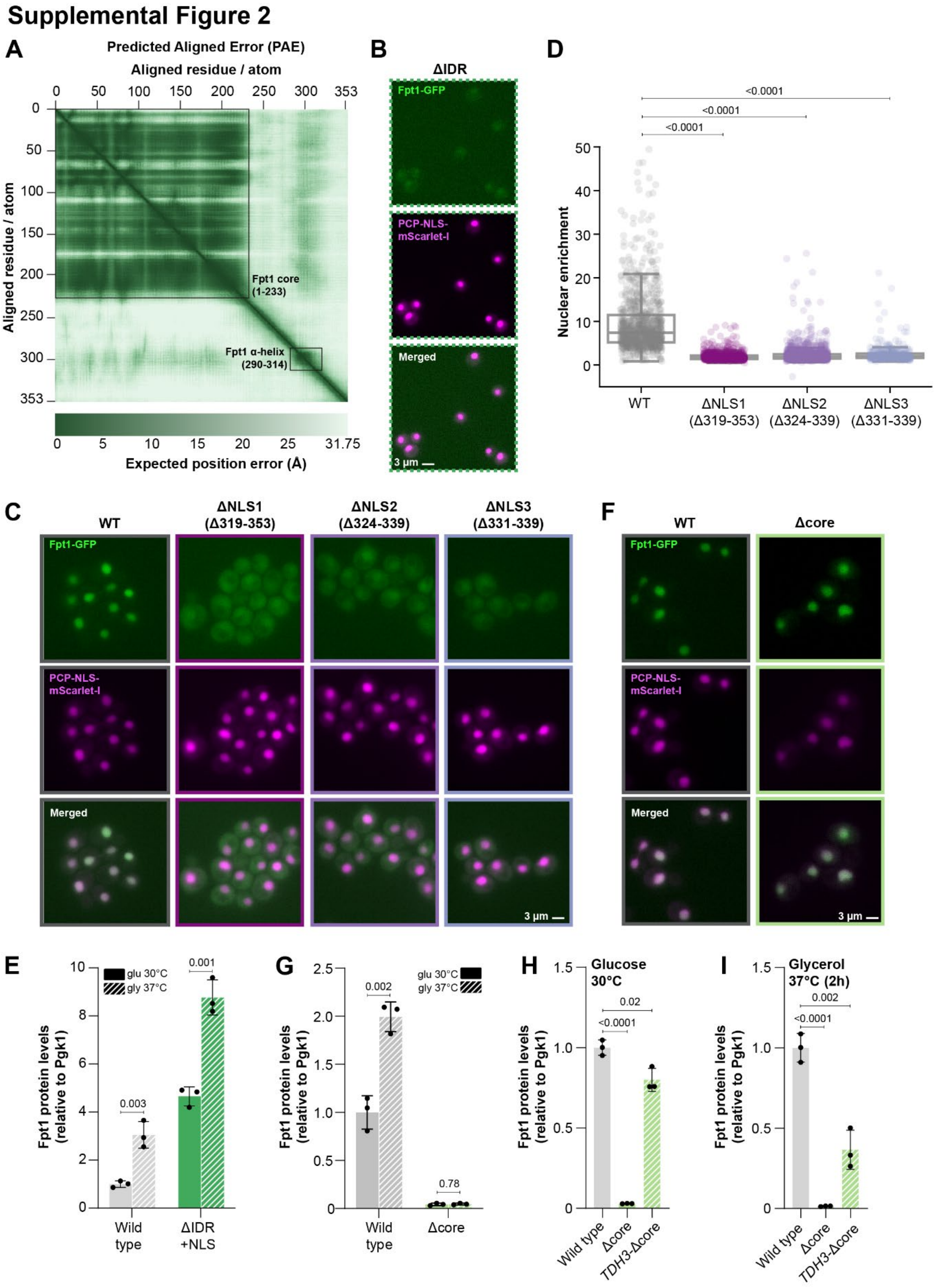

**Supplemental Figure 2. The IDR of Fpt1 encodes a nuclear localization signal and stress-responsive element, while the core domain ensures protein stability.**

a) PAE (Predicted Aligned Error) plot of the Fpt1 AlphaFold2 model.
b) As in Figure 2C but showing higher contrast for Fpt1ΔIDR in the GFP channel.
c) Representative images of wild-type Fpt1 and ΔNLS localization in live yeast cells in glucose 30°C. Several truncation lengths were tested (Fpt1Δ319-353, Fpt1Δ324-339, and Fpt1Δ331-339). From top to bottom: Fpt1-GFP (green), PCP-NLS-mScarlet-I as a nuclear marker (magenta), and the merged channel. Scale bar: 3 µm.
d) Quantification of Fpt1 nuclear enrichment for wild type (n = 968 cells), Fpt1Δ319-353 (n = 805 cells), Fpt1Δ324-339 (n = 1394 cells), and Fpt1Δ331-339 (n = 360 cells). Dots show the data for individual cells; box plots show the distribution of the data where the box indicates the quartiles and whiskers extend to show the distribution, except for outliers. Statistics: bootstrapping (MacKinnon et al.[67])
e) Quantification of Figure 2E.
f) As in (C), representative images of wild-type Fpt1 and Δcore localization in live yeast cells in glucose 30°C.
g) Quantification of Figure 2F.
h) Quantification of Figure 2G.
i) Quantification of Figure 2I.

### Supplemental Figure 3

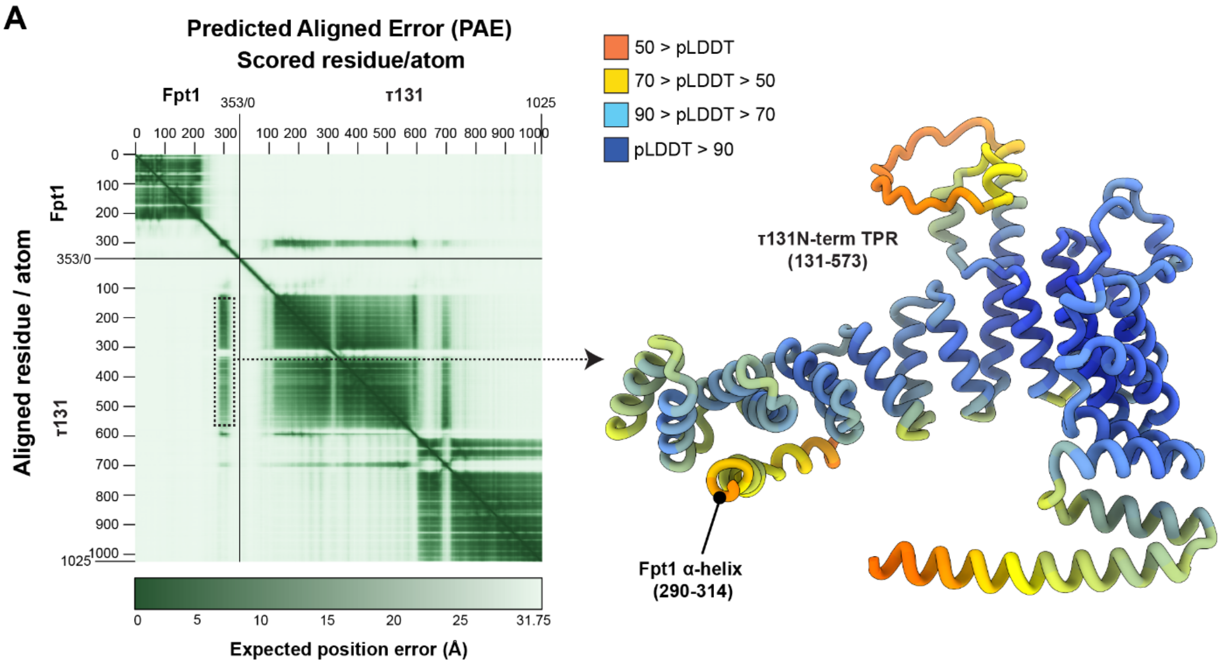


**Supplemental Figure 3. A small region in the disordered region is predicted to interact with τ131 and responsible for the stress response.**

a) Predicted interaction between Fpt1 and τ131. Left: PAE (Predicted Aligned Error) plot of the AlphaFold-Multimer model. The dashed box highlights the region corresponding to the Fpt1–τ131 interface. Right: Predicted structure of the highlighted region, showing the interaction between Fpt1 and the N-terminal TPR domain of τ131. The confidence of the pLDDT (predicted local distance difference test) is shown by different colors.

### Supplemental Figure 4

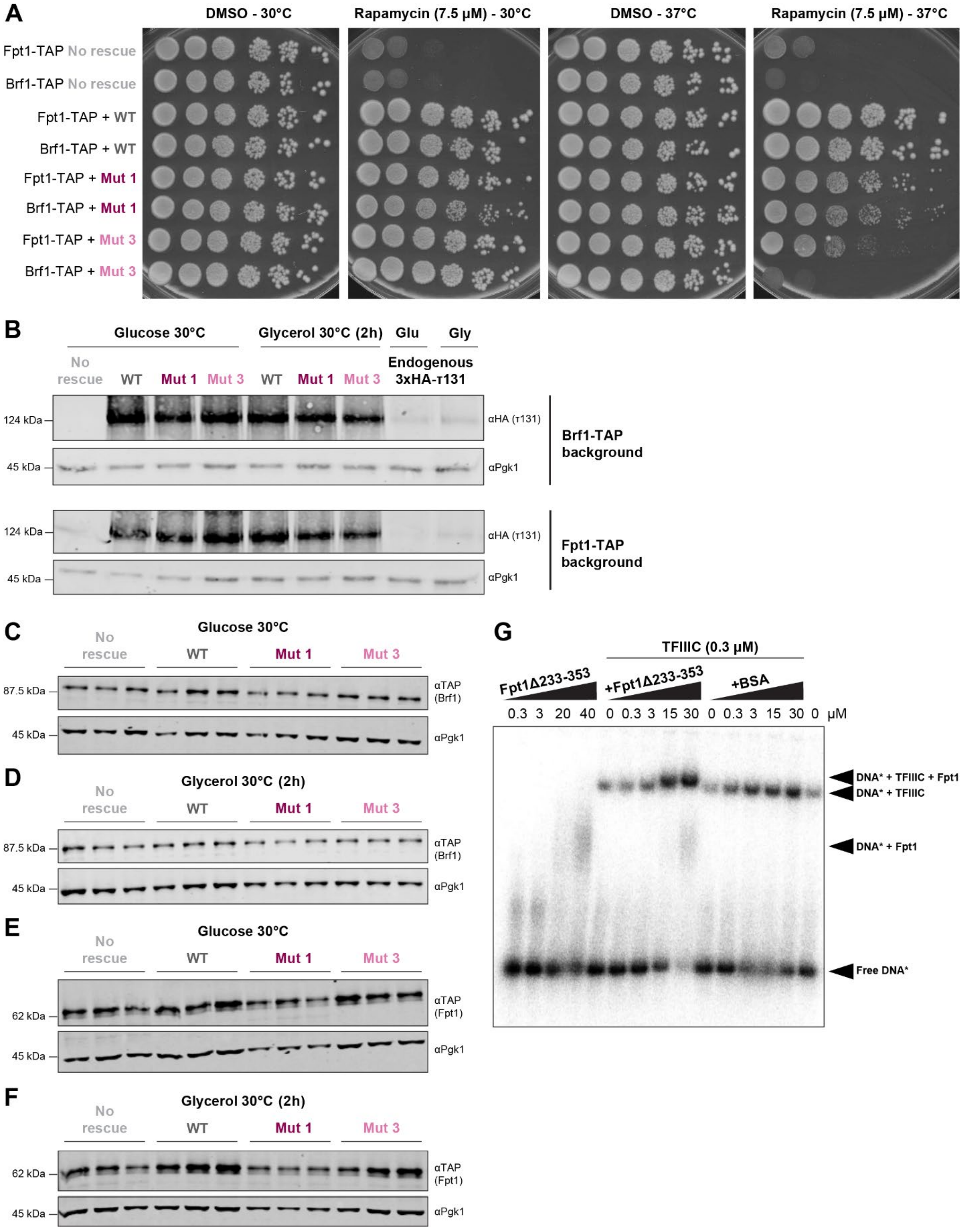

**Supplemental Figure 4. The N-terminal TPR array of τ131 mediates the stress response of Fpt1.**

a) Spot test analysis on rapamycin- (7.5 µM) and DMSO-containing YEP + 2% glucose plates at 30°C and 37°C. No rescue = empty vector without a τ131 copy, WT = wild type τ131 plasmid copy, Mut 1 = τ131 mutant 1, Mut 3 = τ131 mutant 3.
b) Plasmid-copy τ131-3xHA immunoblot (n = 1) for no rescue (empty plasmid), wild type (wild type τ131 plasmid copy), mutant 1 and mutant 3 (mutated τ131 plasmid copies) in a Brf1-TAP (top) and Fpt1-TAP (bottom) background in glucose 30°C and 2h glycerol 30°C. As a control and reference, endogenous τ131-3xHA is displayed.
c) Brf1-TAP immunoblot (n = 3) in a no rescue (empty plasmid), wild type (wild type τ131 plasmid copy), mutant 1 and mutant 3 (mutated τ131 plasmid copies) background in glucose 30°C.
d) As in (C), in 2h glycerol 37°C.
e) As in (C), Fpt1-TAP immunoblot.
f) As in (E), in 2h glycerol 37°C.
g) Electrophoretic mobility shift assay (EMSA) comparing DNA-binding of Fpt1(Δ234-353) to $tRNA^{His}$ promoter DNA in the absence and presence of TFIIIC. DNA* = $tRNA^{His}$ promoter DNA.
